# Human FASTK preferentially binds single-stranded and G-rich RNA

**DOI:** 10.1101/2024.07.16.603671

**Authors:** Daria M. Dawidziak, Dawid A. Dzadz, Mikołaj I. Kuska, Madhuri Kanavalli, Maria M. Klimecka, Matthew Merski, Katarzyna J. Bandyra, Maria W. Górna

## Abstract

Fas-activated serine/threonine kinase (FASTK) is the founding member of the FASTKD protein family, which was shown to regulate the fate of mRNA molecules on multiple levels. The mitochondrial variant of FASTK co-localizes with mitochondrial RNA granules and regulates degradation of mitochondrial mRNAs, whereas the cytoplasmic and nuclear forms of FAST are involved in regulation of alternative splicing, cytoplasmic RNA granule formation and mRNA translation. Despite these multiple roles of FASTK in mRNA biology, the exact rules of RNA recognition by this protein remained undetermined. Here, we demonstrate direct RNA binding by purified human FASTK and show its preference for single-stranded G-rich sites and RNA G-quadruplexes. Addition of FASTK alone was sufficient to achieve protection of mitochondrial mRNAs from degradation by the degradosome. Structural characterization by SAXS showed that FASTK in solution is a monomer with an extended conformation. Point mutagenesis studies supported the structural predictions of an exposed RNA-binding interface in the central helical region, preceded by a smaller, flexibly attached, helical N-terminal domain. We provide the first such extensive *in vitro* characterization of the RNA binding properties for a representative of the FASTKD protein family, and suggest how these intrinsic properties may underly FASTK function in mRNA metabolism.

## Introduction

With six orthologs in humans, the Fas-activated serine/threonine kinase domain-containing (FASTKD) family is one of the largest protein families involved in the regulation of mitochondrially-encoded genes. FASTKD family members often show distinct, or even contradictory influence on certain mitochondrial transcripts and localize to different mitochondrial compartments ^1^. The eponymous FASTK ^2^ was demonstrated to co-localize with mitochondrial RNA granules (MRGs) shown to be associated with post-transcriptional regulation, expression, and degradation of mt-mRNAs ^3–5^. In addition, FASTK has an alternative, non-mitochondrial isoform implicated in the regulation of cell death and autoimmune disorders ^1,2,6–9^. FASTK was first discovered as a partner and a putative kinase of T-cell-restricted intracellular antigen-1 (TIA-1) ^10^, but this activity was questioned since the key residues in its then-proposed active site are not preserved across the family ^11^, leading to replacing the old acronym of the protein (FASTK) with a new alternative name – Fas-activated serine/threonine phosphoprotein (FAST).

FASTKD family members share the same overall architecture: an N-terminal mitochondrial targeting signal (MTS) followed by a region predicted to be mostly helical and containing three conserved sequence domains called FAST_1 (FAST kinase-like protein subdomain 1), FAST_2 (FAST kinase-like protein subdomain 2) and RAP (RNA-binding domain abundant in Apicomplexans) (Figure 1A). The structures of the FASTKD proteins have yet to be experimentally determined, probably due to the presence of disordered regions impeding crystallization. The roles of the FAST_1 and FAST_2 domains remain undetermined, while the RAP domain has been suggested to govern the interactions between FASTKD family members and RNA molecules ^12,13^. The RAP domain comprises ∼60 residues, consists of blocks of charged and aromatic residues ^14^, and shows highly conserved similarities with PD-(D/E)XK nucleases ^15^. It has been suggested that the biological function of FASTKDs relies on the putative catalytic activity of RAP domains ^15^, but so far there has been no experimental confirmation of the nucleolytic activity of FASTKD proteins *in vitro*. Due to predictions suggesting large helical regions in their structures, FASTKD proteins are often linked to pentatricopeptide (PPR) and octatricopeptide repeat (OPR) protein families ^16^. FASTKD proteins, like the PPRs, may function as adaptors, facilitating the identification of specific sequences in mitochondrial transcripts and guiding the interactions with endo- or exonucleases and other RNA-modifying enzymes ^1^.

**Figure 1.**
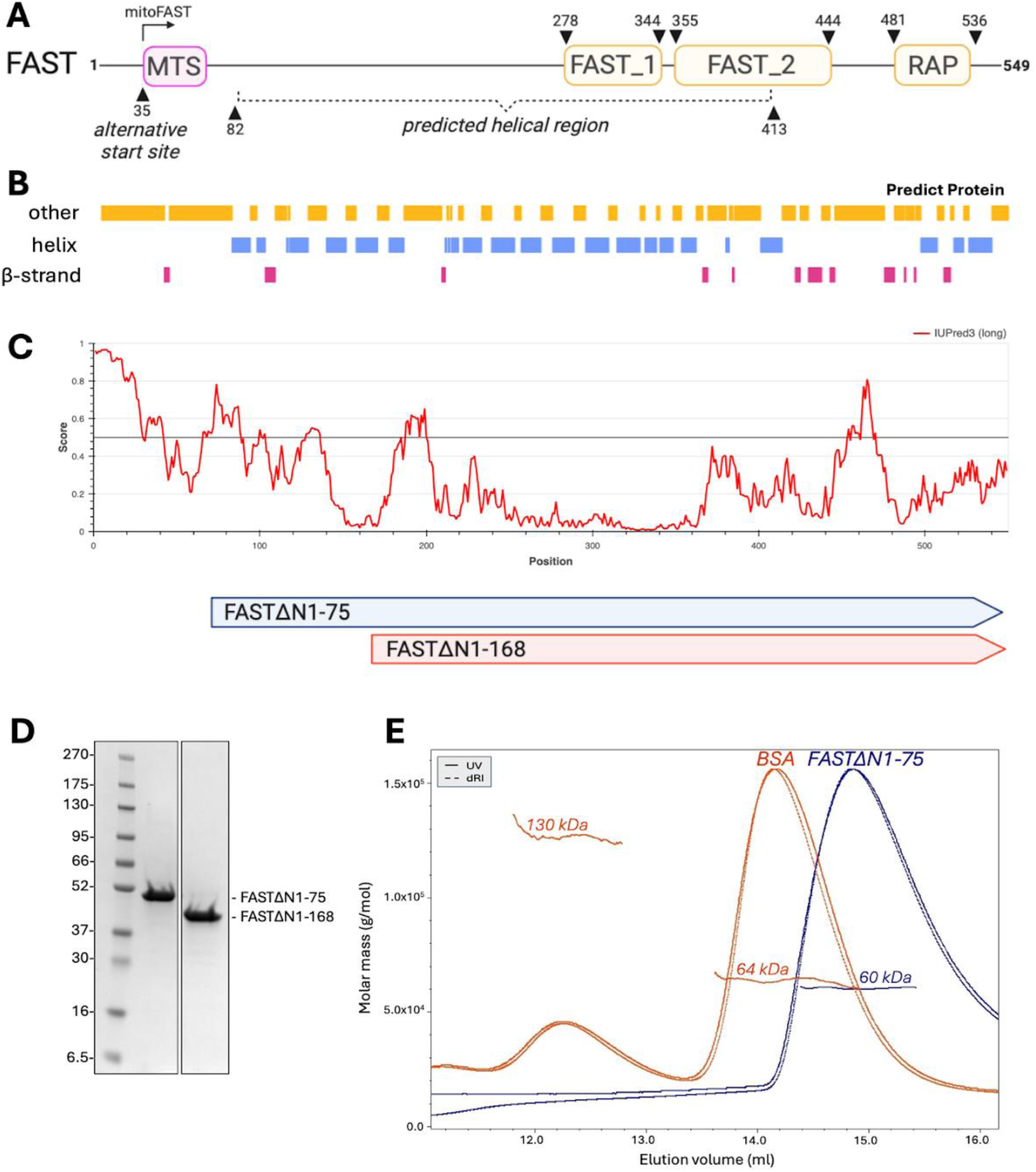
A: Schematic representation of the structure of human FAST protein. FAST_1 (FAST kinase-like protein subdomain 1), FAST_2 (FAST kinase-like protein subdomain 2) and RAP (RNA-binding domain abundant in Apicomplexans) domains are indicated. MTS (mitochondrial targeting signal) indicates the start site of an alternative translational form of FAST residing in mitochondria. B: Diagram showing results of secondary structure predictions done with Predict Protein ^26,27^ – yellow boxes represent regions with unknown secondary structure, purple and pink represent helices and beta-strands, respectively. C: Plot shows results of disordered regions predictions performed with IUPred3 ^28^, where a score between 0 and 1 corresponds to the probability of the given residue being part of a disordered region. Panel below shows amino acids range of FAST constructs -FASTΔN1-75 and FASTΔN1-168. D: SDS-PAGE gel showing samples of two FAST constructs purified according to developed purification procedure. E: SEC-MALS analysis of the purified FASTΔN1-75 (blue trace) and BSA (orange trace). Curves show the UV absorbance and refractive index signal. Horizontal lines show the calculated masses of the eluting components. Domain boundaries were assigned based on the Pfam database in InterPro web suite ^29^. Figure was partially created with BioRender.com.

The maturation of mitochondrial transcripts is executed mostly by removal of flanking tRNAs, according to the tRNA punctuation model, however, there are four transcripts which are not subject to this model^17^. One of them is ND6 mRNA encoding subunit 6 of the NADH dehydrogenase complex (complex I), which exists at steady state as a mature form of 1.0–1.1 kb, along with several precursor RNAs of greater molecular weight referred to as RNA1–3 ^17^. FAST was shown to be particularly important for maturation of this transcript and activity of the whole NADH dehydrogenase complex ^2^. The skeletal and cardiac muscles from mice with disrupted FAST gene displayed ∼60% decrease in NADH dehydrogenase activity. Accordingly, levels of ND6 mRNA were reduced in MEFs isolated from FAST-depleted mice, suggesting that FAST controls expression of this mRNA. Indeed, RNA-IP and NGS revealed that FAST binds both ND6 mRNA and its precursors. Furthermore, it has been shown that human FAST cooperates with the mitochondrial degradosome proteins - polynucleotide phosphorylase PNPase and helicase Suv3p - to generate mature ND6 mRNA by preventing its CDS and 3’UTR from degradation by the degradosome ^2^.

The ND6 gene is the only gene transcribed using the L-strand of the mitochondrial genome, resulting in an mRNA encoded by the H-strand and highly abundant in guanines. Such G-rich L-strand transcripts can theoretically form non-canonical elements called G-quadruplexes (G4s). These structures are created by Hoogsteen base pair interactions which enable formation of a G-tetrad – a planar cycle of four guanines linked by eight hydrogen bonds and stabilized by a monovalent cation, usually a potassium ion ^18,19^. The G-tetrad can be further stabilized by π-π stacking and form a tetrahelical core of the structure. The G4 folding rule states that in order to obtain a G-quadruplex structure, the sequence has to contain four guanine tracts, each composed of at least three guanines and separated by no more than seven other nucleotides (G_3+_N_1-7_G_3+_N_1-7_G_3+_N_1-7_G_3+_N_1-7_) ^20,21^. ND6 has a predicted propensity to form G-quadruplexes ^22^ and such G4-containing RNAs might undergo degradation according to a dedicated surveillance mechanism employing GRSF1, which promotes melting of G4 structures and facilitates degradosome-mediated decay. Interestingly, FAST was shown to co-localize with GRSF1 in mitochondria, but as an inhibitor of the degradosome, FAST is expected to have an opposite effect to GRSF1 on G4-containing RNA. FAST was observed to bind along ND6 with enrichment in certain sections of this transcript ^2^, but the exact sequence motifs or other features of RNA recognised by FAST have not been determined yet.

Numerous studies highlight the role of FAST protein in the execution of mitochondrial cell death pathway ^6–9^. FAST as a regulator of apoptosis was reported to affect alternative splicing of Fas pre-mRNA ^6^, and to boost the expression of mRNAs of anti-apoptotic agents like cIAP and XIAP ^7^, but the potential FAST binding regions in these RNAs remain unknown. FAST is overexpressed in patients with autoimmune diseases including rheumatoid arthritis (RA), systemic lupus erythematosus (SLE), autoimmune diabetes, and multiple sclerosis (MS) ^23,24^. It has been suggested that the overexpression of FAST could be linked to heightened immune system reactivity and the inadequate elimination of immune cells, resulting from their impaired apoptosis. RNA binding properties of FAST could thus play a multipronged role in the cell survival and apoptosis.

Here, we aim to demonstrate and characterize RNA binding by human FAST using highly purified recombinant protein and a range of RNAs. We compared sequences, secondary structures and the nucleotide content of bound RNAs to describe the preference of FAST. Our SAXS-derived experimental structural information supported by machine learning methods provides hints about the RNA-binding interface of FAST.

## Results

### The design and optimization of recombinant FAST purification strategy

The insoluble nature of bacterially expressed FAST has been previously reported and has so far hindered research aiming at solving its 3D structure and explaining the basis of its interactions with RNA molecules ^1^. We relied on secondary structure predictions combined with the Udwary–Merski algorithm (UMA) ^25^, to predict the location of structured and linker regions in the FAST protein sequence (Figures 1B and 1C) and generated two expression constructs (FASTΔN1-75 and FASTΔN1-168) which resulted in highly stable and soluble recombinant proteins (Figure 1D). The purification conditions were optimized, and four crucial modifications were applied in order to increase protein solubility and purity. (i) The purification buffer composition was optimized based on protein melting temperatures using a thermal shift assay in the presence of additives from two commercial screens (Supplementary Figure 1). Buffer pH was adjusted to 8.5 and ammonium sulfate was added throughout to increase protein stability. (ii) 10 mM EDTA was added to the dialysis buffer during the cleavage of His6-SUMO tag, after IMAC HisTrap column purification, to chelate Nickel ions and possibly prevent protein aggregation. (iii) The concentration of NaCl in the lysis buffer was increased to 500 mM and an additional step of polyethyleneimine (PEI) precipitation was added to the procedure to remove nucleic acid contaminations. (iv) An additional step of ion exchange chromatography yielded pure protein freed from other protein contaminants.

The molecular weight and oligomeric state of the recombinant FASTΔN1-75 protein was confirmed using size exclusion chromatography (SEC) coupled with multi angle light scattering (MALS) (Figure 1E). When injected at ∼94 μM concentration, the protein eluted as a single peak with a calculated molecular weight (MW) of 60 kDa, close to that of a monomer (53 kDa as the expected MW from sequence), confirming that FAST construct lacking the N-terminal unstructured region is a monomer in the solution.

FAST was initially described as a putative kinase of TIA-1 ^10^, but this activity has been questioned since the putative kinase active site residues are not preserved across the family ^11^. FASTΔN1-75 kinase activity was assayed *in vitro* in the presence of recombinant TIA1-RRM23 and [γ^32^P]ATP as substrates, but no phosphorylation activity was detected (Supplementary Figure 2). Very slight activity was observed in the gel area of FASTΔN1-75 construct itself, suggesting some residual modification or non-covalent binding between FASTΔN1-75 and [γ^32^P]ATP that might not necessarily be specific. This confirmed that recombinant FASTΔN1-75 does not have protein kinase activity under these conditions, and suggests that regulation of TIA-1 function by FAST is the result of a different mechanism.

### FAST prefers single-stranded and G-rich RNA

In our studies on the RNA-binding specificity of FAST, we first attempted to identify the preferred sequence motifs via SELEX ^30^ using GST-tagged FAST and a random library of short RNA oligonucleotides. This failed to enrich any high-confidence sequence motifs bound by FAST. Due to visible dimerization of amplified RNA oligonucleotides, two rounds of SELEX selection were employed, leading to the identification of three statistically significant RNA sequence motifs (Supplementary Figure 3), two of which happened to be mutually complementary single-stranded RNAs. The biological significance of the discovered motifs was however questionable due to their relatively low abundance (occurring only 680, 227, and 84 times out of approximately 216,000 molecules sequenced). The failure of the SELEX assay is suggestive of the lack of a strong sequence preference for FAST, since in our hands 2-3 SELEX selection rounds were sufficient for another more sequence-specific RNA-binding protein, IFIT2.

To further analyze the RNA targets of FAST, we performed *in vitro* binding assays with synthetic oligonucleotides (Supplementary Table 5) using microscale thermophoresis and native gel electrophoresis. We tested two ssRNAs initially selected by SELEX, a dsRNA formed by these two, and three other structural motifs such as an internal loop and a stem loop (Figure 2A). The comparison of FAST binding measurements (Figure 2B and 2C) showed that the K_D_ values for loop forming, or double-stranded oligonucleotides were notably weaker (155.8 – 237.6 nM) than for single-stranded RNAs (2.7 – 10.3 nM). Moreover, among the single-stranded oligonucleotides, the lowest K_D_ value (2.7 nM) was observed for a G-rich oligonucleotide (5’-GGGUCUGUGGGGUC-3’, henceforth called “G-rich short RNA”) which suggested that FAST might have a propensity for G-rich stretches in single-stranded RNAs. To further examine this hypothesis, we tested modifications of this G-rich oligonucleotide where guanines were replaced with other three nucleotides (Figure 2D). The results indeed emphasized the propensity of FAST for binding G-rich RNA stretches. K_D_ values for A-rich (5’-AAAUCUGUAAAAUC-3’) and U-rich (5’- UUUUCUGUUUUUUC-3’) short RNAs increased by one order of magnitude (78.4 nM and 63.2 nM, respectively), and two orders (212.3 nM) for C-rich short RNA (5’-CCCUCUGUCCCCUC-3’).

**Figure 2.**
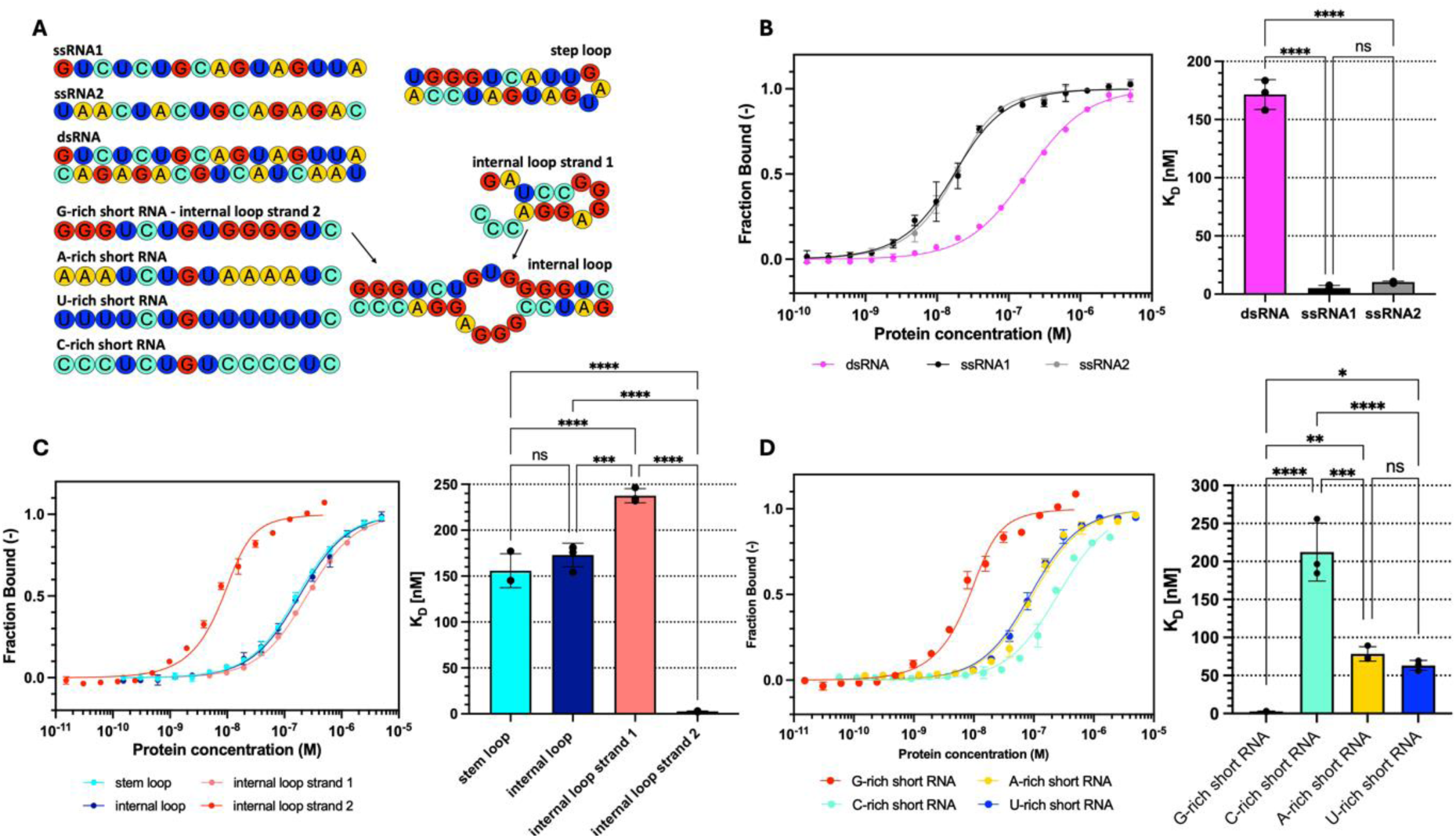
A: Schematic representation of RNA oligonucleotides tested in binding assays. Binding of double-stranded and single-stranded RNA oligonucleotides (B), loop forming RNA oligonucleotides (C) and short RNA oligonucleotides (D) by FASTΔN1-75 was determined by microscale thermophoresis. The binding curves were plotted as a bound fraction of RNA against a protein concentration. Each measurement point was derived from three technical replicates and error bars represent the SD. Statistical significance of pairwise comparisons was calculated with Tukey’s multiple comparisons tests for an ordinary one-way analysis of variance (ANOVA): p > 0.05 (ns); p ≤ 0.05 (*); p ≤ 0.01 (**); p ≤ 0.001 (***); p ≤ 0.0001 (****).

### RNA binding by FAST does not depend on G-quadruplex formation

Our results were consistent with previous reports that FAST binds guanine rich ND6 mRNA and its precursors, which are transcribed from the light strand of the mitochondrial genome ^2^. ND6 mRNA is theoretically prone to form G-quadruplex structures (G4s). We tested FAST for its ability to bind a model RNA G-quadruplex, derived from the telomeric repeat–containing RNA, TERRA (5’UUAGGGUUAGGGUUAGGGUUAGGG-3’) ^31,32^. In order to achieve proper folding of TERRA into its G4 secondary structure, the oligonucleotide was subjected to slow thermal annealing in the presence of potassium ions. The binding assay revealed that FAST indeed binds the TERRA G4 structure with a K_D_ comparable to the one obtained for G-rich short RNA (2.4 nM) (Figure 3A). Along with TERRA we tested two mutants designed to disrupt the formation of the G-quadruplex – the first with point mutations replacing central guanines in guanine triads with adenines and uracils (TERRA AU-rich mutant), and the second with all guanines substituted with cytosines (TERRA C-rich mutant) (Figure 3A). No significant difference in binding was observed for the AU-rich mutant - K_D_ increased less than three-fold (6.4 nM). However, a more pronounced difference in binding resulted from replacing all guanines with cytosines - K_D_ in this case was one order of magnitude higher than for TERRA (32.8 nM). Similarly, the gel shift assay showed a preference towards G-abundant TERRA RNA and visibly lower affinity toward AU- and C-rich TERRA mutant (Supplementary Figure 4).

**Figure 3.**
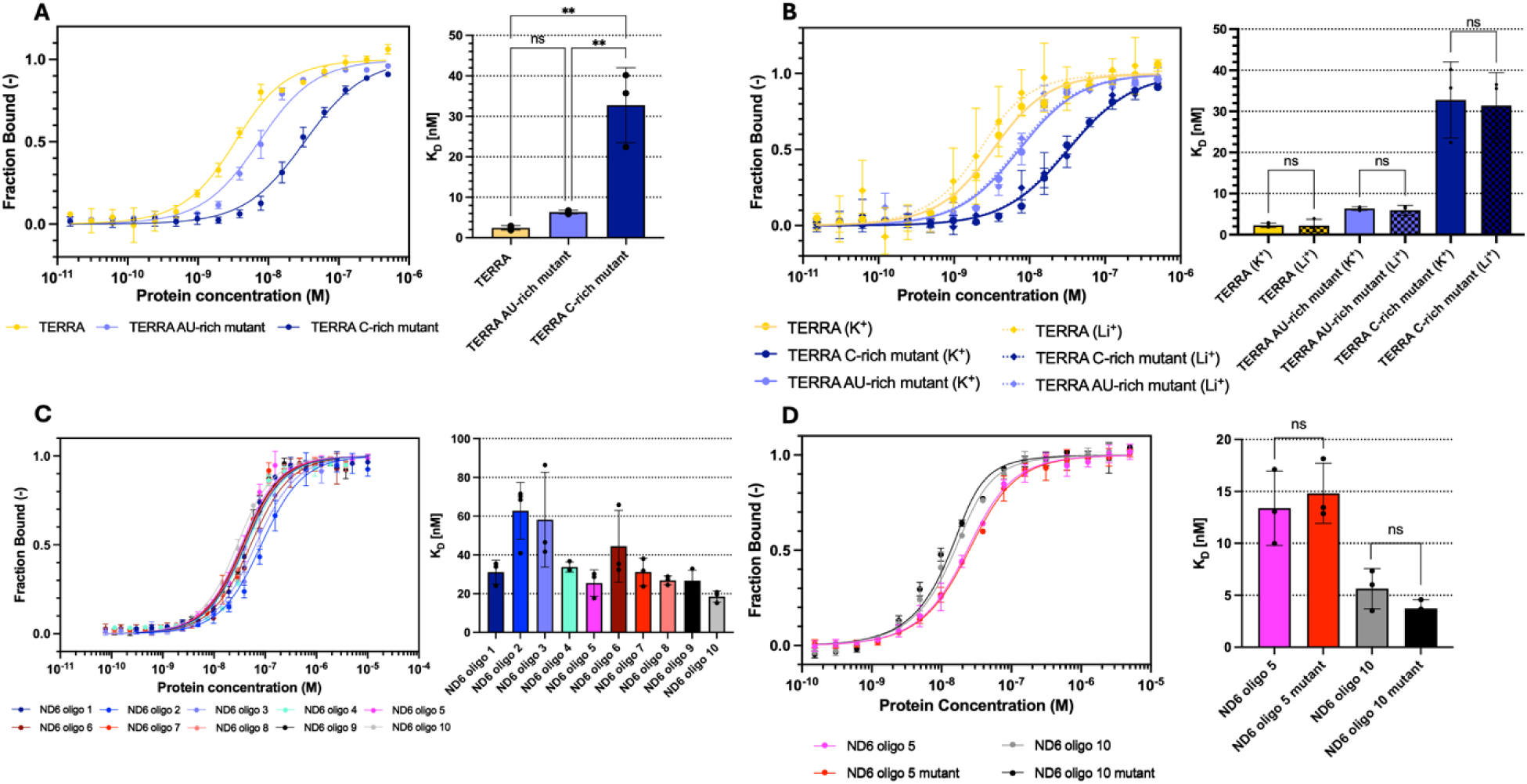
FASTΔN1-75 binding assays performed with microscale thermophoresis. A: Binding of G-quadruplex forming TERRA RNA and its mutants. B: Binding of TERRA RNA and its mutants in the presence of potassium and lithium ions. C: Binding of ten ND6 transcript fragments. D: Binding of RNA oligonucleotides 5 and 10 of ND6 transcript, compared to their binding after introduction of G4-killing mutations. The binding curves were plotted as a bound fraction of RNA against a protein concentration. Each measurement point was derived from three technical replicates and error bars represent the SD. Statistical significance of pairwise comparisons was calculated with Tukey’s multiple comparisons tests for an ordinary one-way analysis of variance (ANOVA): p > 0.05 (ns); p ≤ 0.05 (*); p ≤ 0.01 (**); p ≤ 0.001 (***); p ≤ 0.0001 (****).

To further examine if FAST is capable of binding RNA G-quadruplexes, the same set of TERRA oligonucleotides was subjected to annealing reaction in a regular annealing buffer (50 mM Tris-HCl, 50 mM KCl, 50 mM NaCl; pH 7.5) and a G4-disfavouring buffer containing 50 mM LiCl instead of KCl, to avert the formation of guanine tetrads. Comparison of binding in the presence of K^+^ or Li^+^ proved that the secondary structure of tested oligonucleotides did not have an impact on interactions with FAST. Regardless of the annealing buffer composition, the K_D_ values remained unchanged (Figure 3B).

Consistent results were obtained when we tested the binding of ten ∼100 nt long oligonucleotides derived from the mitochondrial ND6 mRNA (Figure 3C). Interestingly, two of the tested oligonucleotides which exhibited the lowest K_D_ values in FAST binding assays (oligo 5 - 25.5 nM, oligo 10 - 18.5 nM) comprised G-rich sequences capable of forming G4s structures. In order to assess the relevance of these structures in FAST-RNA interactions, oligonucleotides were modified to disrupt the formation of putative G-quadruplexes by introduction of adenines and uracils between the guanine nucleotides (Supplementary Table 6). Such mutations had no impact on interactions with FAST (Figure 3D), which might suggest that FAST binds G-rich stretches of single stranded RNAs that might be separated with adenines and uracils, and which do not necessarily have to constitute a G-quadruplex structure.

The influence of the G4-disrupting mutations on the secondary structure of all the tested oligonucleotides was confirmed by detection of G-quadruplexes with the Thioflavin T dye (Supplementary Figure 5). The intensities of the fluorescence emission spectra measured for TERRA and ND6 oligonucleotides mutants were visibly weaker than for the original RNAs, suggesting that the introduced mutations indeed decreased formation of G-quadruplexes.

We attempted to characterize the complex of FAST with the model TERRA RNA by SEC-MALS analysis. Based on the SEC-MALS data, we concluded that FAST exists as a monomer in the solution, but we could not exclude the possibility of formation of higher order oligomeric states upon binding to RNA, nor binding of more than one TERRA molecule to a single FAST molecule. However, when FASTΔN1-75 was mixed at a 1:1 molar ratio with TERRA RNA, the resulting complex was insoluble and could not be observed by SEC-MALS. Regardless of whether the protein was mixed with RNA at a high concentration, or first mixed at a low concentration and then concentrated as a complex – this mixture had a tendency to precipitate from the solution. After centrifugation of such complex, only a tiny peak was observed by SEC-MALS for the FAST monomer, suggesting that FAST forms higher order protein-RNA assemblies with propensity to aggregation upon binding to RNA.

### FAST protects bound RNA from degradation by the degradosome

We next tested FAST binding to several full-length, *in vitro* transcribed, mitochondrial mRNAs (Figure 4A). Surprisingly, all these transcripts bound very well regardless of their calculated G % or predicted G4 content (Supplementary Table 7), with COX2 mRNA having the strongest binding affinity. We selected ND6 and COX2 mRNAs for degradation assays to investigate the effect of FAST binding on the downstream functional consequences.

**Figure 4.**
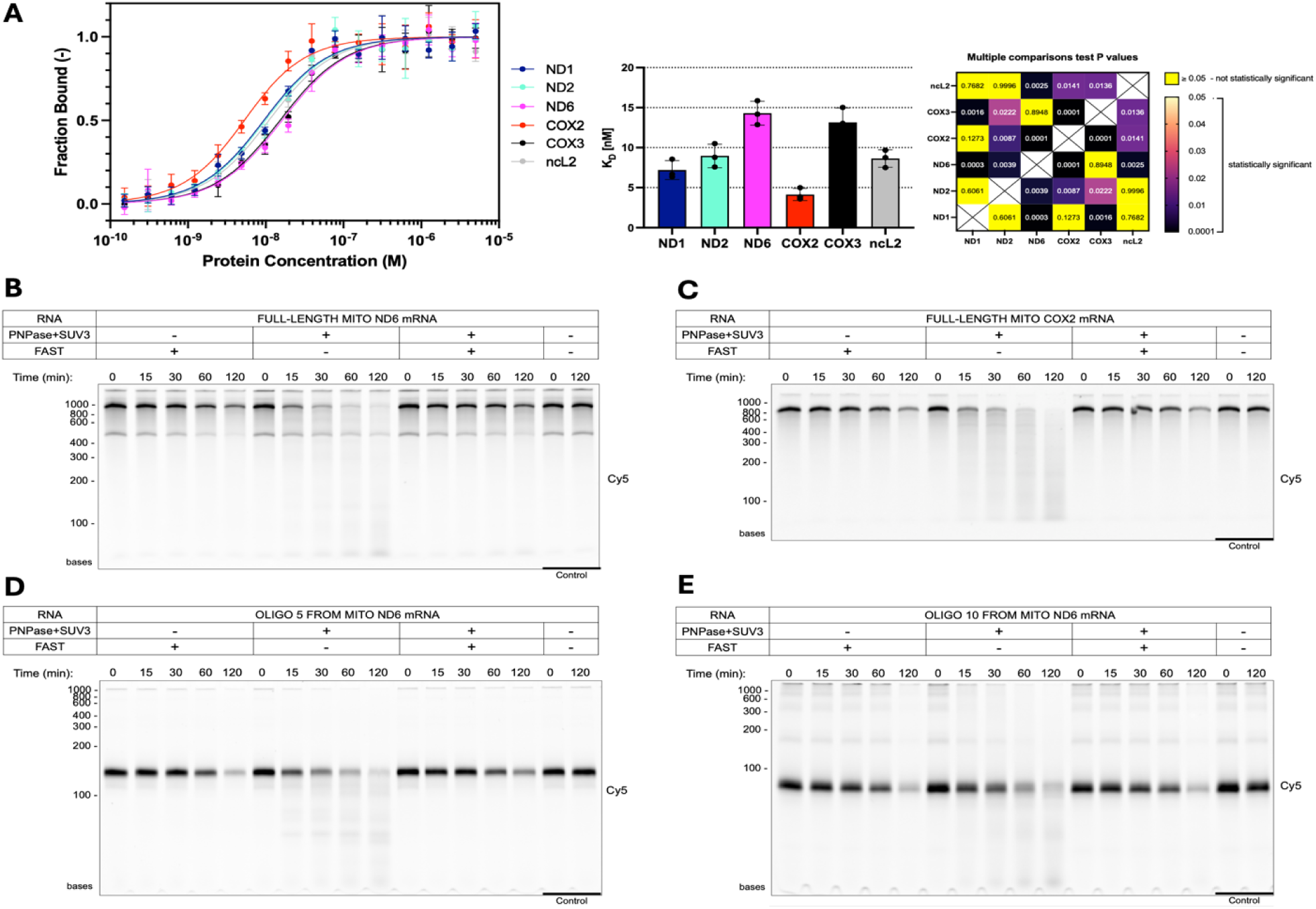
A: Binding of six mitochondrial transcripts by FASTΔN1-75 protein, determined by microscale thermophoresis. The binding curves were plotted as a bound fraction of RNA against a protein concentration. Each measurement point was derived from three technical replicates and error bars represent the SD. Mean K_D_ values of individual experiments with SD are shown. The p-values of Tukey’s multiple comparisons tests for ANOVA comparing mean K_D_ values are presented as a heatmap. A significant result is p < 0.05. X indicates no valid comparison. In vitro degradation of Cy5-body-labeled ND6 mRNA (B), COX2 mRNA (C), and ND6 mRNA fragments (oligo 5 - D, oligo 10 - E) by recombinant PNPase and Suv3p proteins and the impact of FAST on the RNA degradation. Visualized is the Cy5 signal after denaturing PAGE. Reaction samples were collected at 0, 15, 30, 60, 120 min timepoints. The two last lanes present RNA after incubation without addition of recombinant proteins.

Previously, cell-line studies using RNA-IP and NGS revealed that FAST binds ND6 mRNA and its precursors and cooperates with the mitochondrial degradosome to generate mature ND6, by preventing its CDS and 3’UTR degradation ^2^. To test the role of *in vitro* FAST activity in RNA protection, we set up an RNA protection assay containing bacterially expressed and purified components of the mitochondrial degradosome complex (the 3’-5’ exoribonuclease polynucleotide phosphorylase PNPase and the helicase Suv3p), FASTΔN1-75 and *in vitro* transcribed Cy5-labeled mitochondrial mRNAs. The outcome of our RNA protection assays confirmed the observations made formerly by *in vivo* assays. All tested RNA substrates were degraded by the degradosome, however in the presence of FAST the degradation activity was notably diminished (Figure 4B-E). In accordance with the RNA binding preference studies carried on full-length mitochondrial transcripts, the protection activity seemed to be independent of the nucleotide composition of the used substrates. Both the G-rich ND6 and C-rich COX2 mRNAs were equally protected in reactions containing recombinant FAST. This protective effect also extended to the shorter ND6 oligonucleotides 5 and 10.

We observed some decrease in the concentration of the full-length transcripts with the addition of FAST to the reactions with prolonged incubation times, and we investigated the possibility that this effect might be attributed to residual FAST nuclease activity or a nuclease contamination. However, this decrease in full-length RNAs was not associated with any intermediate degradation products derived from the body-labeled substrates, unlike the clear ladder visible in all degradosome-treated samples. Mass spectrometry analysis of two independently produced batches of our recombinant FAST did not indicate any abundant nuclease contaminants (Supplementary Table 9). Moreover, we have already observed solubility problems with FAST-RNA mixtures. In line with these observations, we thus speculated that the most likely cause for RNA loss might be that RNA-FAST complexes precipitate during the prolonged incubation of reaction mixtures.

### A low-resolution structural model of FAST reveals a potential RNA binding interface

The optimized expression and purification procedure of two FAST protein constructs resulted in excellent, homogenous samples suitable for structural studies. Although FAST eluded crystallization, we obtained the first experimental structural information for FAST from SEC-SAXS measurements (Supplementary Table 8). We calculated the electron density distribution and compared the reconstructed structural envelopes ^33^ for FASTΔN1-75 (Figure 5) and FASTΔN1-168 (Supplementary Figure 6) with structural predictions generated by various machine learning methods. To validate our low-resolution experimental densities, we fitted FAST atomic models created with RoseTTAFold ^34^ or AlphaFold2 ^35^. The atomic models fit the experimental data well in the C-terminal region of the protein, comprising the conserved FASTKD protein family domains - FAST_1, FAST_2 and RAP. The alignment of the two models suggested that a flexible proline-rich linker (amino acids 188 to 196) might allow the protein to adopt both an open and a closed conformation. While the RoseTTAFold model displayed an open conformation with the N-terminal helical domain distant from the C-terminal RAP domain, the AlphaFold2 model presented a more compact conformation with the helical domain angled towards the C-terminal region (Figure 5A). AlphaFold3 model of FASTΔN1-75 assumed an even more closed conformation of the N-terminal helical domain, which further suggests its possibly mobile nature.

**Figure 5.**
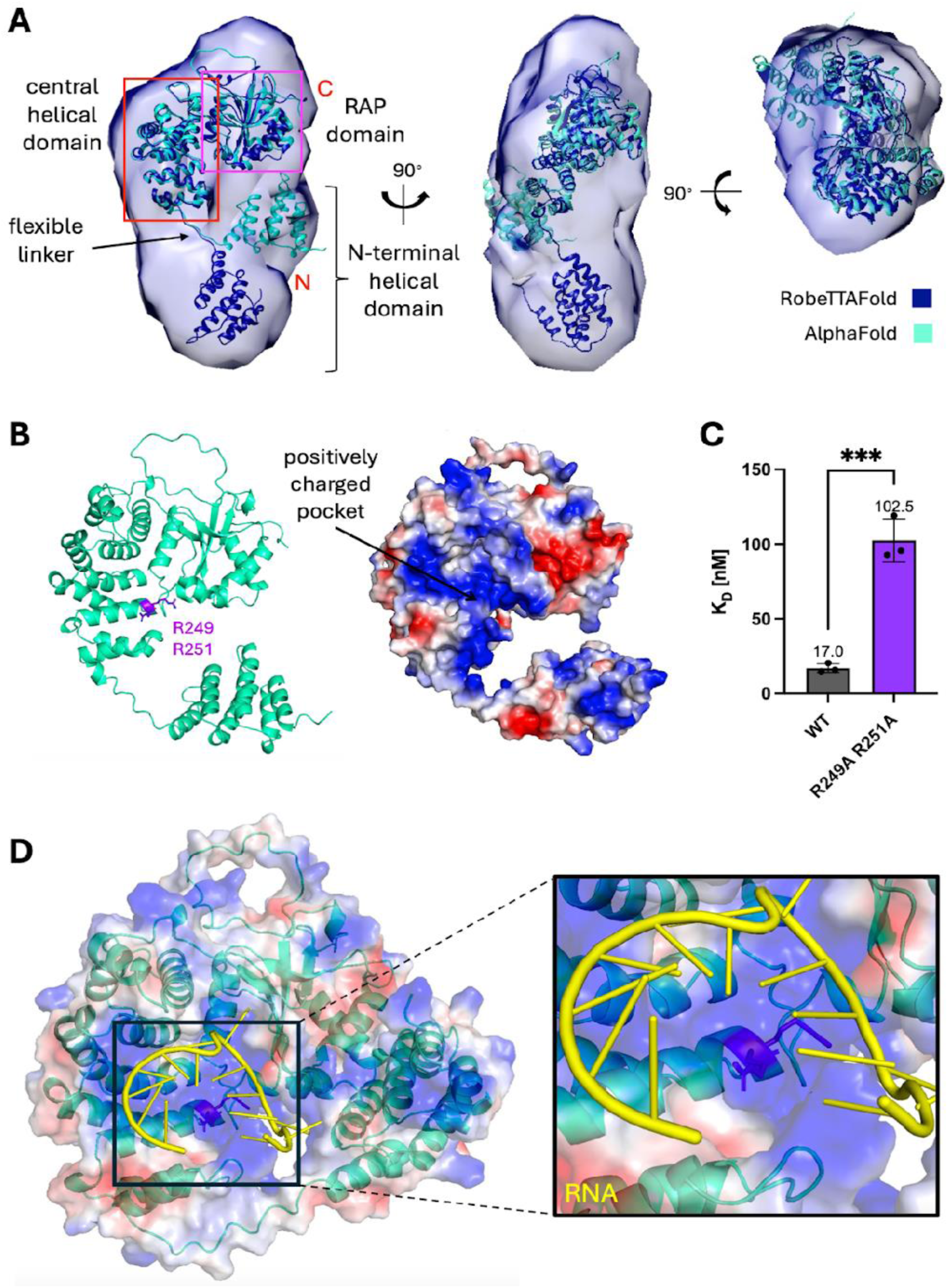
A: Front, side, and top views of FASTΔN1-75 prediction models fitted in the electron density generated by DENSS algorithm ^33^ based on the SEC-SAXS measurements. Model created with RobeTTAFold ^34^ (dark blue) was fitted into the map using the Fit in Map tool in UCSF Chimera software, then the AlphaFold2 ^35^ model (aquamarine) was aligned to it using the MatchMaker comparison tool. B: Map of protein contact potential generated for AlphaFold2 prediction model of FASTΔN1-75 allowed to discover positively charged pocket, putatively involved in RNA binding. C: Mutations in residues R249 and R251 significantly lowered FASTΔN1-75 ability to bind fragment 5 of mitochondrial ND6 RNA. D: The AlphaFold3 ^36^ prediction of FASTΔN1-75-RNA complex structure utilizes RNA binding site encompassing residues R249 and R251.

The analysis of the surface electrostatic potential of FASTΔN1-75 models revealed a positively charged region within the central helical domain of the protein – a putative RNA binding site (Figure 5B). To probe the localization of the putative RNA binding interface and the accuracy of the predicted model, we mutated several batches of positively charged residues, and characterized RNA binding by the mutant FASTΔN1-75. Residues embedded within the central pocket (R249, R251, K275, K279 and R286), alongside other positively charged, exposed amino acids (Supplementary Figure 7) were changed to alanines or methionines to neutralize their charges. We purified mutants and evaluated their binding of the ND6 oligo 5. Mutations at residues R249 and R251 decreased RNA binding 5-fold when compared to the wild-type protein (Figure 5C), and mutations at the nearby K275, K279 and R286 were also markedly effective (Supplementary Figure 7B), whereas other mutants had no direct effects (and in two cases caused reduced protein solubility). Finally, AlphaFold3 ^36^ which allows facile prediction of protein-RNA complexes also placed ND6 oligo 5 in the vicinity of the R249-R251 region (Figure 5D).

## Discussion

Direct RNA binding by human FAST has been suggested by evidence from human cell line studies and backed by the involvement of its homologues in RNA metabolism in other species. Here, we provided the first direct evidence of RNA binding by FAST, due to the availability of highly purified and homogenous samples of the recombinant protein. We characterized RNA binding preference of FAST in terms of sequence specificity, RNA structure and nucleotide content. Our *in vitro* binding and degradation assays showed that purified FAST is able and sufficient to recapitulate previous *in vivo* reports.

Our success in FAST purification might be partially due to the careful removal of the bound (bacterial) RNA, since we observed decreased solubility of FAST-RNA mixtures. Purified FAST was monomeric, but in our hands precipitated upon the addition of RNA, which suggests formation of higher-order protein-RNA assemblies (condensates), reminiscent of a phase transition. This is in line with the descriptions of FAST as a component of RNA granules both in the mitochondria and in the cytoplasm. The likely flexible nature of FAST has so far rendered it poorly suitable for high-resolution studies, but our low-resolution SAXS data agree with most of the current machine learning structural predictions. The SAXS data suggest an extended, open conformation of FAST in solution, whereas the closed conformation predominated in Alphafold predictions – the experimental data overcome a known caveat of the current state of the machine learning methods which cannot fully recapitulate protein dynamics such as the positioning of the mobile parts of the protein. Our mutagenesis study provided evidence for the localization of the putative RNA-binding interface in the central region of FAST; it is interesting to speculate that FAST might “clamp” around RNA molecules using the R249/R251-lined “saddle” and the flexibly-connected N-terminal domain.

We were able to delve into the RNA-binding properties of FAST using the most amenable short oligonucleotides. We failed to detect any RNA sequence-specific features, but FAST was sensitive to the nucleotide content of RNA, with preference towards G-rich oligonucleotides and discrimination against C-rich ones. This propensity for G-rich oligonucleotides was affected by the strand pairing, with ssRNA being preferred over dsRNA or stem loops. FAST binding to RNA was relatively insensitive to the introduction of single adenine or uracil nucleotides in the G-tracts, as well as to the formation of G-quadruplexes. This suggests FAST has permissive RNA binding properties. Interestingly, in a small screen of six full-length mitochondrial mRNAs, all were bound well by FAST regardless of their G- or G4-content. This suggested that longer transcripts might perhaps benefit from cooperativity or a local concentration effect to promote coating of the RNA by FAST. Notably, FAST binding to COX2 mRNA was tighter than to the ND6 mRNA, the best-described natural target of FAST. *In vitro* folded, isolated mRNAs most likely differ from their physiologically-relevant structures and conformations found *in vivo*, and their accessibility in the cell is expected to be further regulated by proteins, cellular machineries, membranes and localization. Similarly, in case of FAST binding to RNA, the preferences of this protein demonstrated here using shorter oligonucleotides will be compounded by many additional factors in the cell, especially those regulating RNA accessibility or FAST localization. While the intrinsic properties of FAST might direct it to binding G-rich transcripts such as ND6, we expect other factors to increase selectivity of FAST *in vivo*.

We demonstrated that addition of FAST alone was sufficient to compete with and inhibit degradation by the degradosome, which recapitulates well the known protective effect of FAST on ND6 mRNA. This mechanism seems to correlate with direct RNA binding by FAST, as it was also observed with other substrates. FAST binding selectivity might thus drive the protective effect of FAST. In the simplest approximation, RNA binding by FAST may result in a physical obstacle due to FAST-RNA interactions outcompeting degradosome processivity (*i.e.* the degradosome is deterred since it is unable to displace FAST “markers” deposited on G-rich sites or G-quadruplexes (Figure 6)). However, liquid-liquid phase separation might be yet another mechanism through which FAST may regulate RNA localization and accessibility *in vivo*. These downstream effects are not easily disambiguated *in vitro* nor *in vivo*, as single FAST-RNA complexes are difficult to obtain for study but future single-molecule studies in cells could help with this issue.

**Figure 6.**
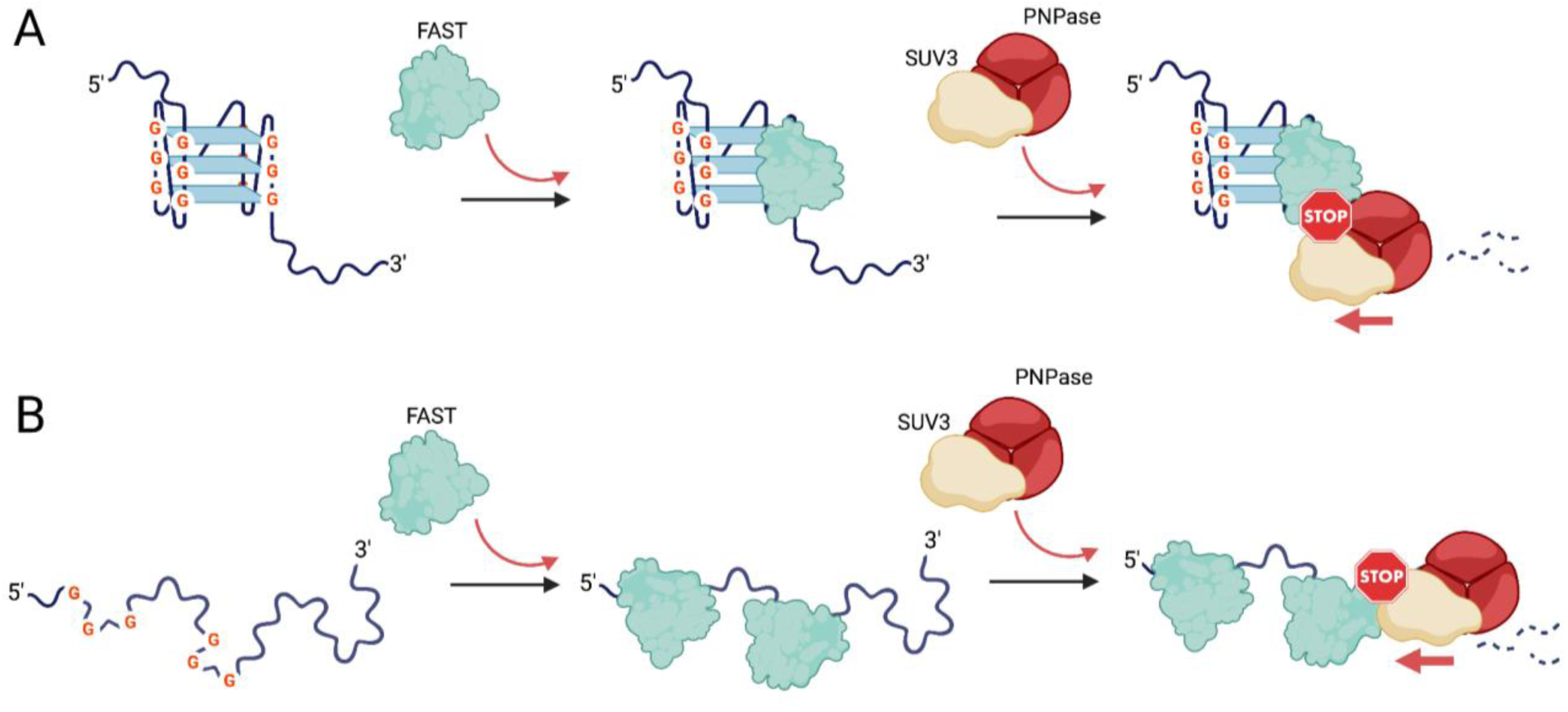
FAST protects mRNA from the degradation by the degradosome by preventing its access to G-rich sites (A) and G-quadruplexes (B).

Our observations explain how FAST might regulate the degradation of ND6 and other G-rich transcript fragments in the mitochondria, but the question arises how the same binding properties of FAST might be used in other cellular compartments which lack the degradosome. In the nucleus, FAST might bind G-rich sequences in precursor or mature mRNAs and regulate their splicing or stability through interplay with other proteins. Although locally G-enriched regions in transcripts are ubiquitous and therefore their comprehensive analysis may be difficult, specific G4-forming regions have started to be characterized for their role in RNA stability and accessibility ^37^. G4s are highly enriched at splice site regions and are preferably formed within the non-template strand of the genome, leading to their formation on the pre-mRNA ^38–42^. Moreover, G4s are proposed to play a role in stress granule formation via their interactions with RBPs ^43–46^, which correlates with the reports of FAST as a scaffolding protein mediating transitions between stress granules and processing bodies ^47^. Indeed, mRNA known to be regulated by FAST contain sequences predicted to form G4s - for example, the 5’UTRs of cIAP ^48^ and Bcl-2 mRNAs ^49^ - and these G-rich regions might serve to recruit FAST. Our results suggest that the role of G-rich RNA recognition by FAST and its role in the nuclear RNA metabolism and cytoplasmic RNA granules should be the subject of future studies.

## Methods

### Cloning of plasmids for bacterial expression

The full list of vectors and DNA starters used in the study is provided in the Supplementary Tables 1 and 2. The expression vector pCA528, containing a sequence of His6-SUMO tag, was a kind gift from Prof. Owen Pornillos (University of Virginia). The vector used for the expression of Suv3p protein was described previously ^22^. The pGEX-4T1 vector used for cloning of GST-FASTΔN1-168 for the SELEX assay was a kind gift from Dr. Martyna Nowacka (University of Warsaw). Vectors pDONR223-FASTK and pET28a_TIA1 were purchased from Addgene (#23712 and #106095).

All constructs for the bacterial expression of proteins (pCA528-FASTΔN1-75, pCA528-FASTΔN1-168, pCA528-TIA1-RRM23 and pGEX-GST-FASTΔN1-168) were obtained by SLIC cloning ^50,51^. FAST encoding DNA was amplified from pDONR223-FASTK plasmid and was cloned into pCA528 vector, followed by transformation to *E. coli* Top10 and kanamycin selection. The mutagenesis of pCA528-FASTΔN1-75 was performed by PCR amplification of whole plasmid DNA using a suitable pair of primers introducing intended mutations (Supplementary Table 2). Amplification was followed by plasmid circularization by simultaneous phosphorylation and ligation of plasmid ends, and transformation of the *E. coli* Top10 and kanamycin selection. For cloning of pCA528-TIA1-RRM23, DNA of TIA-1 protein (aa 80-292) was amplified from pET28a-TIA1 plasmid and was cloned into pCA528 vector, followed by transformation to *E. coli* Top10 and kanamycin selection. For cloning of pGEX-GST-FASTΔN1-168, DNA of FAST ORF was amplified from pDONR223-FASTK plasmid and was cloned into pGEX-4T1 vector, followed by transformation to *E. coli* Top10 and ampicillin selection.

### Cloning and *in vitro* transcription of mitochondrial mRNA

All RNA oligonucleotides up to 25 nucleotides long were purchased from either Merck or Future Synthesis. The pCR™-Blunt II-TOPO™ vector was a kind gift from Dr. Marcin Ziemniak (University of Warsaw). The mitochondrial DNA isolated from HeLa cells was a kind gift from Dr. hab. Roman Szczęsny (Institute of Biochemistry and Biophysics, Polish Academy of Sciences). Human mitochondrial *MT-ND6* gene was amplified from mitochondrial DNA and inserted into pCR™-Blunt II-TOPO™ vector for further *in vitro* transcription. The sequence of *MT-ND6* gene was amplified by PCR reaction and used in TOPO™ cloning reaction according to the manufacturer’s instructions. For *in vitro* transcription of fragments of ND6 mRNA bearing mutations removing possible formation of G-quadruplexes, the pCR™-Blunt II-TOPO™-ND6 plasmid was mutated using procedure analogous to the one described above.

pCR™-Blunt II-TOPO™-ND6 vector was used as a template for *in vitro* transcription of full length ND6 mitochondrial transcript, its ∼100 nt long fragments and mutants. The selected DNA sequences were amplified by PCR reaction with forward starters containing overhangs with sequence of T7 RNA polymerase promoter (5’-TAATACGACTCACTATAGG-3’). The list of starters used in the experiment is specified in Supplementary Table 3. 20 μl *in vitro* transcription reactions were prepared on ice with the following ingredients: DNA template (300 - 1000 ng), 1x T7 RNA polymerase buffer, rNTPs (5 mM each*), 1 mM DTT, 3% DMSO, 0.25 μl RiboLock RNAse inhibitor (Thermo Scientific), 2 μl home-made T7 RNA polymerase. * For transcription reaction of fluorescently labeled RNAs, 0.5 mM Cy5-CTP or Cy5-UTP was included in the reaction and the concentration of corresponding unlabeled ribonucleotide was lowered to 3.5 mM. Reactions were incubated at 37°C for 4 h and transferred on ice. The template DNA was then digested by addition of 1U of TURBO DNAse (Thermo Scientific) and incubation at 37°C for another 30 min. The reactions were quenched by addition of 50 mM EDTA. The products of the transcription were then purified with QIAGEN RNeasy Mini Kit according to the manufacturer’s instructions and visualized by RNA PAGE.

*In vitro* transcriptions of additional mitochondrial transcripts (*ncL2*, *COX2*, *COX3, ND1* and *ND2*) were performed using pRS147X vectors, which were a kind gift from dr hab. Roman Szczęsny. pRS147X vectors have a built-in sequence of T7 RNA polymerase promoter, a sequence of each transcript and restriction enzymes sites, enabling adjustment of the length of transcript’s poly(A) tail. The *in vitro* transcription reactions were prepared as described above and products were analyzed by RNA-PAGE.

### RNA annealing

The RNA annealing procedure was performed in 1x Annealing Buffer (50 mM Tris-HCl pH 7.5, 50 mM KCl, 50 mM NaCl) with the use of thermal cycler and according to the following steps: (i) 5 min denaturation at 95°C, (ii) slow cooling to RNA’s T_M_, (iii) 30 min incubation at RNA’s T_M_ and (iv) slow cooling to 4°C. The whole procedure was spread over time to take 2 h. RNAs were then transferred on ice and immediately used for further experiments.

### Thioflavin T assay

The Thioflavin T reactions were prepared by mixing 2 μM Thioflavin T (Thermo Scientific), 2 μM annealed RNA in 1x Annealing Buffer. Reactions were then transferred to 96-well plates (PS, F-bottom/chimney well) (Greiner) and measured by Tecan M200 Pro Plate Reader with excitation at 430 nm and the fluorescence emission scan between 460 and 700 nm every 2 nm.

### Protein expression

The overexpression of FASTΔN1-75, FASTΔN1-168 (Supplementary Table 4), GST-FASTΔN1-168, TIA1- RRM23, PNPase and Suv3p proteins was performed in *E. coli* BL21-CodonPlus(DE3)-RIL cells with the induction of protein expression at OD_600_ of 0.5-0.8 with 1 mM IPTG, followed by overnight incubation at 20°C.

### Purification of recombinant FAST proteins

Harvested cell pellets were suspended in Lysis Buffer (150 mM Tris-HCl pH 8.5, 5 % (v/v) glycerol, 500 mM NaCl, 200 mM ammonium sulfate, 10 mM arginine, 10 mM glutamic acid, 5 mM β-Me), freshly supplemented with 1 mM PMSF, 10 mM MgCl_2_, 1 mg/mL hen egg lysozyme and 10 μg/mL DNase I, and incubated for 1h at 4°C on a rocking platform. Cells were disrupted by sonication (7 min, 2s on/2s off, 60% amplitude) on ice. The lysate was centrifuged at 20000 xg, 4°C for 30 min and the obtained supernatant was subjected to polyethyleneimine (PEI) precipitation of nucleic acid contaminations as described in ^52^ : the lysate supernatant was placed in a glass beaker on a magnetic stirrer at 4°C and 9% PEI solution was added dropwise to the supernatant, until 0.1% concentration was reached. The solution was left with stirring for 20 min and then centrifuged at 20000 xg, 4°C for 20 min.

The supernatant was then filtered through a 40 μm syringe filter and applied on HisTrap HP column (Cytiva) attached to a Bio-Rad FPLC system and equilibrated with HisTrap Buffer A (50 mM Tris-HCl pH 8.5, 5 % (v/v) glycerol, 50 mM NaCl, 50 mM ammonium sulfate, 10 mM arginine, 10 mM glutamic acid, 5 mM β-Me). The recombinant His6-SUMO tagged protein was then eluted with increasing gradient of HisTrap Buffer B (1M Imidazole, 50 mM Tris-HCl pH 8.5, 5 % (v/v) glycerol, 50 mM NaCl, 50 mM ammonium sulfate, 10 mM arginine, 10 mM glutamic acid, 5 mM β-Me). Fractions containing protein of interest were combined and dialyzed overnight at 4°C with 50 ml of home-made recombinant SUMO protease (1 mg/ml), against Dialysis Buffer (50 mM Tris-HCl pH 8.0, 5 % (v/v) glycerol, 50 mM NaCl, 150 mM ammonium sulfate, 10 mM arginine, 10 mM glutamic acid, 10 mM EDTA, 10 mM imidazole, 5 mM β-Me). The cleaved protein was then diluted 2x in HiTrap SP Buffer A (50 mM Tris-HCl pH 8.0, 5 % (v/v) glycerol, 50 mM ammonium sulfate, 10 mM arginine, 10 mM glutamic acid, 5 mM β-Me) to decrease the NaCl concentration and facilitate protein binding to the column. The solution was then applied on HiTrap SP column (Cytiva) and the recombinant protein was eluted with increasing gradient of HiTrap SP Buffer B (50 mM Tris-HCl pH 8.0, 5 % (v/v) glycerol, 1000 mM NaCl, 50 mM ammonium sulfate, 10 mM arginine, 10 mM glutamic acid, 5 mM β-Me). Eluted protein fractions were then concentrated to volume < 4 ml using Amicon^®^ Centrifugal Filter Units (10 kDa MWCO) by centrifugation at 3000-5000 xg, 4°C for 10-15 min. Concentrated protein was centrifuged to remove all precipitates at 10000 xg, 4°C for 15 min and applied on HiLoad 16/600 Superdex 75 pg column (Cytiva), equilibrated in Storage Buffer (50 mM Tris-HCl pH 8.5, 5 % (v/v) glycerol, 50 mM NaCl, 250 mM ammonium sulfate, 10 mM arginine, 10 mM glutamic acid, 1 mM DTT). The fractions containing clean protein were pooled, concentrated as described above, flash frozen in liquid nitrogen and kept at −80°C until used. All significant samples obtained during expression and purification were analyzed by SDS-PAGE.

### Thermal shift assay

Purified FASTΔN1-75 at a concentration of 1 mg/ml, initially in buffer containing 50 mM Tris-HCl pH 7.5, 5 % (v/v) glycerol, 400 mM NaCl, 10 mM arginine, 10 mM glutamic acid and 0.5 mM TCEP, was mixed with 5x concentrated SYPRO™ Orange Protein Gel Stain (Invitrogen) and 10x diluted components of Additive Screen (Hampton Research) and Solubility Screen (Jena Bioscience) in 25 μl reaction volume. Reactions were prepared on 384-well PCR plates, sealed with optically clear sheet of adhesive, briefly centrifuged, and subjected to thermal gradient produced by a CFX384 Touch Real-Time PCR Detection System (Bio-Rad).

### Microscale thermophoresis

Purified FASTΔN1-75 protein was dialyzed against MST Buffer (1x PBS, 5% (v/v) glycerol, 1 mM DTT) at 4°C for 2h and centrifuged to remove all precipitates at 10000 xg, 4°C for 10 min. The concentration of protein was measured using NanoDrop spectrophotometer and then protein was diluted in MST buffer to a concentration of 2 μM. Samples of annealed Cy5 labeled RNA oligonucleotides were diluted in MST buffer supplemented with 0.05% Tween-20 to concentration of 40 nM. Sixteen binding mixtures were prepared by 1:1 mixing of an RNA solution (constant concentration), with protein solution (a 2-fold dilution series). Reaction mixtures were pre-incubated for 5 min on ice, centrifuged for 2 min and loaded into capillaries. The measurements were performed using a Monolith NT.115 device (Nanotemper Technologies) at a constant temperature of 22°C, according to the manufacturer’s instructions. Each measurement point (mean ± SD) was derived from three technical replicates. The fitting of the dose-response curves to a one-site binding model was performed using MO Affinity Analysis v.2.3 software, supplied by the device manufacturer. The fit curves were then exported and plotted as a bound fraction of RNA against a protein concentration using Prism 10 (GraphPad). The mean and SD of the K_D_ values obtained from three separate measurements were calculated and then the statistical significance of pairwise comparisons of K_D_ values was calculated using two methods: (i) ordinary one-way analysis of variance (ANOVA) test with Tukey’s multiple comparisons test or (ii) the two-tailed unpaired parametric t-test, where p > 0.05 (ns); p ≤ 0.05 (*); p ≤ 0.01 (**); p ≤ 0.001 (***); p ≤ 0.0001 (****).

### Electrophoretic mobility shift assay (EMSA)

Solutions of FASTΔN1-75 protein and RNA oligonucleotides were prepared as described above for MST measurements. The binding mixtures were prepared by mixing an RNA solution of constant concentration, with a series of 2x dilutions of protein solution in the concentration range from 1000 to ∼8 nM. Additionally, two control reactions were included – one containing protein alone at the highest concentration used, and the second control with RNA alone. Reactions were then incubated for 5 min on ice, mixed with 10x EMSA loading dye (30% (w/v) Ficoll 400 (Sigma-Aldrich) in water, supplemented with Orange G dye (Leica)) and separated by native gel electrophoresis in TG buffer using 4–20% Mini-PROTEAN^®^ TGX Stain-Free™ gels (Bio-Rad), subjected to 30 min pre-run. Gels were resolved at 100V until ready and visualized with a ChemiDoc Imaging System. The intensities of RNA bands were quantified using ImageLab software.

### Analysis of oligomeric state by SEC-MALS

Purified FASTΔN1-75 protein was dialyzed against SEC-MALS Buffer (1x PBS, 50 mM ammonium sulfate, 5% (v/v) glycerol, 1 mM DTT) for 2h at 4°C. The analysis was carried out by injecting 100 μL of purified protein at a concentration of 5 mg/ml and onto a Superdex 200 Increase 10/300 GL column (Cytiva) equilibrated with SEC-MALS Buffer and connected to NGC Scout 10 FPLC System (Bio-Rad). A solution of bovine serum albumin in SEC-MALS Buffer at the same concentration and volume was used as a calibration standard. Particle size and refractive index data were collected with a RefractoMax 520 Refractive Index Detector (ERC Inc) equipped with an in-line miniDAWN TREOS static light scattering detector (Wyatt Technologies). Data were recorded and processed using ASTRA software (Wyatt Technology) using a single dn/dc value of 0.185 mL/g ^53^.

### Analysis of recombinant FASTΔN1-75 by mass spectrometry

Proteins were precipitated by methanol/chloroform precipitation and suspended in 100 mM HEPES, pH 8.0 containing 5 mM TCEP and 10 mM CAA (2-chloroacetamide). Samples were incubated with 0.4 μg trypsin overnight in a thermo-shaker set to 37°C. The digestion was terminated by adding trifluoroacetic acid (TFA) to 1% final concentration, and digested peptides were desalted with a C18-StageTip. Prior to LC-MS measurement, the peptide fractions were resuspended in 0.1% TFA, 2% acetonitrile in water. Chromatographic separation was performed on an Easy-Spray Acclaim PepMap column (50 cm long × 75 µm, Thermo Fisher Scientific) at 55 °C by applying 60 min acetonitrile gradients in 0.1% aqueous formic acid at a flow rate of 300 nl/min. An UltiMate 3000 nano-LC system was coupled to a Q Exactive HF-X mass spectrometer (data-dependent mode, resolution 30,000, m/z 200) via an easy-spray source (all Thermo Fisher Scientific). Up to 15 of the most abundant isotope patterns with charges 2-5 from the survey scan were selected with an isolation window of 1.3 m/z and fragmented by higher-energy collision dissociation (HCD) with normalized collision energies of 27, while the dynamic exclusion was set to 30 s. The maximum ion injection times for the survey scan and the MS/MS scans (acquired with a resolution of 15,000 at m/z 200) were 45 and 22 ms, respectively. The ion target value for MS was set to 3 × 10^6^ and for MS/MS to 10^5^, and the intensity threshold for MS/MS was set to 2×10^4^.

### Kinase activity assays

TIA1-RRM23 protein purification procedure was performed as described in ^54^, with minor modifications. [γ^32^P]ATP (Hartmann Analytic), control protein kinase (SnRK2.6) and myelin basic protein (MBP) were a kind gift from Prof. Grażyna Dobrowolska (Institute of Biochemistry and Biophysics, Polish Academy of Sciences). The putative kinase activity of FAST protein was analyzed *in vitro* in the presence of recombinant FASTΔN1-75 and recombinant TIA1-RRM23 and [γ^32^P]ATP as substrates. The protein components of the reactions were combined in Kinase Assay Buffer (25 mM Tris-HCl pH 7.5, 5 mM EGTA, 1 mM DTT, 30 mM MgCl_2_). Reactions were then initiated by addition of 50 mM ATP supplemented with 1 µCi of [γ^32^P]ATP and were immediately placed at 30 °C. After 40 min the reactions were quenched by addition of 5x SDS-PAGE Sample Loading Buffer (50 mM Tris-HCl pH 6.8, 2% (w/v) SDS, 10% (v/v) glycerol, 100 mM β-Me, 0.01% Bromophenol Blue) and boiling for 5 min. 15 μl of each sample was separated by SDS-PAGE at 200V, and phosphorylated substrates were visualized by autoradiography.

### Systematic Evolution of Ligands by Exponential enrichment (SELEX)

GST-FASTΔN1-168 protein was expressed in 100 ml of transformed BL21(DE3) RIL *E.coli* culture, induced with 1mM IPTG and cultured at 18°C overnight. The harvested pellet was resuspended in 1xPBS supplemented with 1mM PMSF, cells were lysed by sonication (3 min, 30s on/30s off, 40% amplitude) and centrifuged for 45 min at 17 000 xg, 4°C. Centrifuged lysate was then mixed with 400 ml of Glutathione Sepharose beads (Cytiva) (washed with miliQ water and 1xPBS) and incubated for 1h at 4°C with slow rotation. Beads were then washed 3 times with 1xPBS, and protein was eluted in 2 portions (1 and 0.5 mL each) by incubation in GST Elution Buffer (50 mM Tris-HCl pH 8.5, 5 % glycerol, 50 mM NaCl, 200 mM ammonium sulfate, 1 mM DTT, 10 mM reduced glutathione) with shaking at 1000 rpm, 4°C for 15 min on a thermomixer. Eluted protein was decanted from the beads, flash frozen and stored at −80°C.

80 picomoles of purified GST-FASTΔN1-168 were incubated for 1h at 4°C in 1 ml of SELEX Binding Buffer (20 mM Tris-HCl pH 7.5, 0.1 % Triton X-100, 100 mM NaCl, 1 mM DTT) with 30 ml of Glutathione Sepharose beads. After incubation beads were washed, resuspended in 1 ml of SELEX Binding Buffer supplemented with 2 µg/ml of poly dIdC (Sigma-Aldrich) and mixed with 13 μg of *in vitro* transcribed library of random RNA oligonucleotides. The binding reaction was incubated with rotation for 1h at 4°C. Beads were then washed with SELEX Washing Buffer (20 mM Tris-HCl pH 7.5, 0.1 % Triton X-100, 250 mM NaCl, 1 mM DTT), and bound RNA oligonucleotides were released by proteolysis of FAST protein in 400 μl of SELEX Washing Buffer supplemented with 25 μg of Proteinase K (Invitrogen) for 1h at 37°C, 800 rpm with shaking. The eluted RNA was precipitated in 1 ml of ice cold 100% ethanol with the addition of 3 μl of linear acrylamide (Invitrogen) and stored overnight at −20°C. The next day, RNA was pelleted by centrifugation at 4°C, 15000 rpm for 30 minutes, washed with cold 80% ethanol and dried by incubation at 37°C. The dried pellet was then resuspended in water and reverse-transcribed with 1st Strand cDNA Synthesis Kit for RT-PCR (AMV) (Roche) following the manufacturer’s protocol. The obtained cDNA was then *in vitro* transcribed according to the procedure for T7 RNA polymerase and used in the subsequent selection reaction. After the second iteration of selection, the amplified cDNA was sequenced by Next Generation Sequencing with Genome Sequencer Illumina NovaSeq 6000 PE100 (the Genomics Core Facility, Centre of New Technologies, University of Warsaw). The obtained data were analyzed using the motif discovery tool MEME (Multiple Em for Motif Elicitation) ^55,56^.

### RNA protection assays

The purification of Suv3p was performed according to the FAST protein purification procedure. The only modification was introduced during the lysis of the pellets, when cells were homogenized using the PandaPLUS homogenizer. The recombinant human PNPase protein was prepared using a modified protocol for PNPase purification described in ^57^. Degradation assay reactions were carried at 37°C in 50 μl reaction volume containing 50 or 200 nM RNA and indicated protein/s (FASTΔN1-75, Suv3p or PNPase) at concentrations of 1 μM each in RNA degradation buffer (20 mM Tris-HCl pH 7.5, 50 mM NaCl, 50 mM KCl, 1 mM MgCl_2_, 2 mM Sodium phosphate) freshly supplemented with 1 mM DTT, 1 mM ATP (Thermo Fisher Scientific) and 1U/⌠l RiboLock RNAse inhibitor (Thermo Fisher Scientific). Reactions were started by the addition of RNA previously subjected to slow annealing and placing at 37°C. Full length mitochondrial RNA transcripts were used in the assay at the concentration of 50 nM, while shorter oligonucleotides (12 - 125 nt) were used at the concentration of 200 nM. At selected time points 10 μl portions of reactions were mixed with 10 μl of 0.5 mg/ml Proteinase K in Proteinase K buffer (100 mM Tris-HCl pH 7.5, 150 mM NaCl, 25 mM EDTA, 1% (w/v) SDS) and were immediately placed at 50°C for 20 min. Samples were then removed from the thermo block, mixed with 10 μl of 2x formamide RNA loading buffer (95 % (v/v) Formamide, 1 mM EDTA, 0.01% (w/v) Orange G dye (Leica), in deionized water) and flash frozen in liquid nitrogen. When all samples were collected, they were heated for 5 min at 95 °C, briefly centrifuged and then the entire sample was applied and separated on a Urea PAGE gel in 1x TBE buffer. To enable good separation and visualization of short products of degradation, Urea PAGE was performed on 20 x 20 cm gels prepared using the UreaGel System from National Diagnostics. The gels were then stained by incubation with SYBR Gold Stain for 20 min in TBE buffer or were directly scanned with Cy5 channel on ChemiDoc (BioRad) system. The intensities of RNA bands were quantified using ImageLab software.

### Size-Exclusion Chromatography Coupled with Small-Angle X-ray Scattering (SEC-SAXS)

x-ray scattering measurements of FASTΔN1-75 and FASTΔN1-168 were conducted at the B21 beamline of Diamond Light Source (Didcot, UK) ^58^ using 13.1 keV X-rays at 50 × 50 μm beam-size at detector with 0.8 × 2 mm photon cross-section at sample and flux 4 x 1012 photons/s, and approximately 620 frames (exposure time = 3 s, q range from 0.0045 to 0.34 Å^−1^) were collected per sample using Eiger 4M detector. Protein samples were separated by the in-line SEC system (Agilent 1260 HPLC) connected to Superdex 200 Increase 3.2/300 column (GE Healthcare) equilibrated in FAST SAXS Buffer (50 mM Tris-HCl pH 8.5, 5% (v/v) glycerol, 200 mM ammonium sulfate, 50 mM NaCl, 0.5 mM TCEP, 10 mM glutamic acid, 10 mM arginine) using a flow rate 0.075 mL/min. SEC-SAXS data processing was performed using the BioXTAS RAW 2.2.1 software package ^59,60^ with extension launching ATSAS 3.2.1 software suite ^61^ using RAW as GUI. The SEC-SAXS run images were plotted as scattergram (integrated scattering intensity vs. frame number). Buffer regions were selected automatically using liquid chromatography (LC) series analysis tool and sample regions were selected based on the flatness of Rg parameter. The averaged profiles from buffer subtracted images were used for the Guinier analysis. The Guinier fit for FASTΔN1-168 data was performed automatically while FASTΔN1-75 data required truncation of Guinier region to reduce the deviation in the distribution of residuals. The molecular weight of samples was calculated using following methods: (i) from volume of correlation ^62^, (ii) from Porod volume ^63^, (iii) by Shape&Size ^64^ and (iv) from Bayesian inference method (a part of ATSAS package) ^65^. P(r) functions were generated using BIFT algorithm ^66^. In both datasets the qmax values were truncated to 0.1604 in order to obtain reasonable Dmax values and uniform distribution of normalized residuals. The reconstruction of electron density was performed using DENSS program ^33^ with default settings. All visualizations of protein structures and ED maps were prepared in UCSF Chimera v. 1.15 and PyMOL v. 2.5.5.

## Acknowledgements

We would like to thank Dr Jan Kutner, Aleksandra Jurska and Dr Anna Laskowska for help in setting up the early stages of the project and their technical assistance. We thank Prof. Andrzej Dziembowski and Dr Roman Szczęsny for helpful discussions. We thank Prof. Krzysztof Woźniak and the Core Facility for Crystallography and Biophysics for hosting, training and infrastructure for this work. We thank Dr Nikul Khunti for SAXS data collection and Dr Marcin Ziemniak for consultations on the SEC-SAXS data processing.

## Funding

This work was supported by the National Science Centre, Poland [grant agreement 2014/15/D/NZ1/00968 to MWG, 2020/37/N/NZ1/02348 to DMD, 2020/37/K/NZ1/02312 to KJB] and EMBO Installation Grant [3315]. The research leading to these results has received funding from the Norwegian Financial Mechanism 2014-2021 2020/37/K/NZ1/02312. Core Facility for Crystallography and Biophysics was supported by the Foundation for Polish Science under the European Regional Development Fund, TEAM TECH Core Facility [POIR.04.04.00-00-31DF/17]. NGS was performed thanks to Genomics Core Facility CeNT UW (RRID:SCR_022718), using NovaSeq 6000 platform financed by Polish Ministry of Science and Higher Education [decision no. 6817/IA/SP/2018 of 2018-04-10].

## Author contributions

DMD and MWG designed the study; DMD, KJB and MWG wrote the first draft of the manuscript; DMD, DAD, MIK and MK purified proteins; DMD, DAD and MIK performed RNA binding assays; DMD and MIK performed cloning and mutagenesis; DMD performed structural studies and RNA degradation assays; DMD and MMK performed kinase assay; DMD and KJB analyzed degradation assays; MM, DMD and MWG analyzed structural predictions.

## Data availability

SAXS data were deposited in SASBDB under accession entry identifiers SASDVV2 and SASDVW2.

